# Decision and response monitoring during working memory are sequentially represented in the human insula

**DOI:** 10.1101/2022.10.25.513764

**Authors:** A. Llorens, L. Bellier, A.O. Blenkmann, J. Ivanovic, P.G. Larsson, J.J. Lin, T. Endestad, A.-K. Solbakk, R.T. Knight

## Abstract

Emerging research supports a role of the insula in human cognition. Here, we used intracranial EEG to investigate the spatiotemporal dynamics in the insula during a verbal working memory (vWM) task. We found robust effects for theta, beta, and high frequency activity (HFA) during probe presentation requiring a decision. Theta and beta band activity showed differential involvement across left and right insulae while sequential HFA modulations were observed along the anteroposterior axis. HFA in anterior insula tracked decision making and subsequent HFA was observed in posterior insula after the behavioral response. Our results provide electrophysiological evidence of engagement of different insula subregions in both decision-making and response monitoring during vWM and expand our knowledge of the role of the insula in complex human behavior.

## Introduction

Interest in the human insula has increased commensurate with the development of improved methods to study this structure^1,2^. The insula is located deep within the lateral sulcus of the brain^3^. It is divided by the central insular sulcus into anterior (three short gyri and one accessory gyrus) and posterior parts (two long gyri), and outlined by the circular sulcus^4,5^. The insula shares bidirectional connections with multiple regions of the neocortex, including prefrontal cortex (PFC), anterior cingulate cortex (ACC), supplementary motor area (SMA), parietal, occipital, and temporal cortices. It also connects with medial temporal and adjacent limbic structures^6,7^ as well as with the basal ganglia^8,9^. This extensive connectivity places the insula as an ideal neuroanatomical hub supporting numerous cognitive and limbic functions^1,10–12^ with overlapping and distinct roles played by its different subregions and across both hemispheres^13–15^. Some reports note differential inter-hemispheric involvement, with the left insula preferentially observed in response inhibition and language processing and the right insula in error prediction, error detection, memory, and attention^6,16–22^.

Cytoarchitectural, imaging, and modeling studies report a spatial posterior-to-anterior gradient linked to a progressive integration of complex behaviors including top–down control^17,23–26^. Broadly, the insula can be divided into three anatomical subregions which play distinct functional roles in human cognition, albeit with some overlap^21,27^. The posterior insula is preferentially connected to parietal, occipital, and temporal association cortices and is involved in interoceptive experiences, somatosensory, viscerosensory and sensorimotor functions, speech production, and primary and association auditory responses^5,17,23,24,28–31^. The anterior insula, composed of a ventral and a dorsal part^32^, receives inputs from frontotemporal regions such as inferior frontal gyrus (IFG), dorsolateral PFC (DLPFC), ACC, amygdala, limbic areas, and frontopolar regions. The ventral anterior insula is involved in emotion and empathy processing^13,21,26,33^. The dorsal anterior plays a key role in core cognitive functions including executive control^22,34^, decision making, attention, salience processing through its connection with the ACC^35^, task switching, and error and deviancy detection^1,2,14,17,19,23,24,36–38^. Neuroimaging studies have also reported anterior insula involvement in working memory (WM^39,40^,see^13^ for a review) and verbal WM (vWM) processes such as maintenance, interference resolution, and recency effects^41–48^.

High-order processes such as WM, which require a decision and a subsequent motor response, might engage both anterior and posterior insulae, yet the spatiotemporal dynamics of the insula during WM has not been explored.

Here, we investigated the involvement of the insula during vWM using intracranial electrophysiology (iEEG). We analyzed different frequency band modulations recorded from electrodes implanted in different subregions of the insula of patients undergoing pre-surgical invasive monitoring for potential treatment of drug-resistant epilepsy^49^. We focused on theta and beta band oscillations as these are known to underpin different aspects of the (v)WM process^50,51^ and have been reported along the anterior–posterior axis of the insula^52^. We also extracted high frequency activity (HFA), known to be an index of local neuronal population activity^53–56^ which has been observed in cognitive processes such as WM^57^. Lastly, we assessed the relationship of insular frequency band activity and behavioral performance.

## Results

### Behavioral results

Twelve patients performed a vWM task requiring them to encode and maintain in memory a list of five successive letters, then to answer whether a probe letter had been presented in the list (Figure 1A). They executed the task well with an average of 79.2% of correct answers (range 56% - 95.1%). The mean response time (RT) across patients was 1031.8ms (SD 173.6ms; Figure 1B). No significant correlation was found between RT and accuracy across patients (r=−.38, p=.24). The age of participants was not correlated with RT (p=.75) nor with the percentage of correct answers (p=.70). No gender effect was found on RT (p=.55) or percentage of correct answers (p=.43).Correct trials for positive (i.e., probe letter presented in the current list) and negative (i.e., probe letter not presented in the current list) response conditions did not significantly differ in number (total number of correct trials across patients: 654 vs. 694) nor in RT (mean RT of correct vs. incorrect trials across patients: 1036ms vs. 1030.7ms).

**Figure 1:**
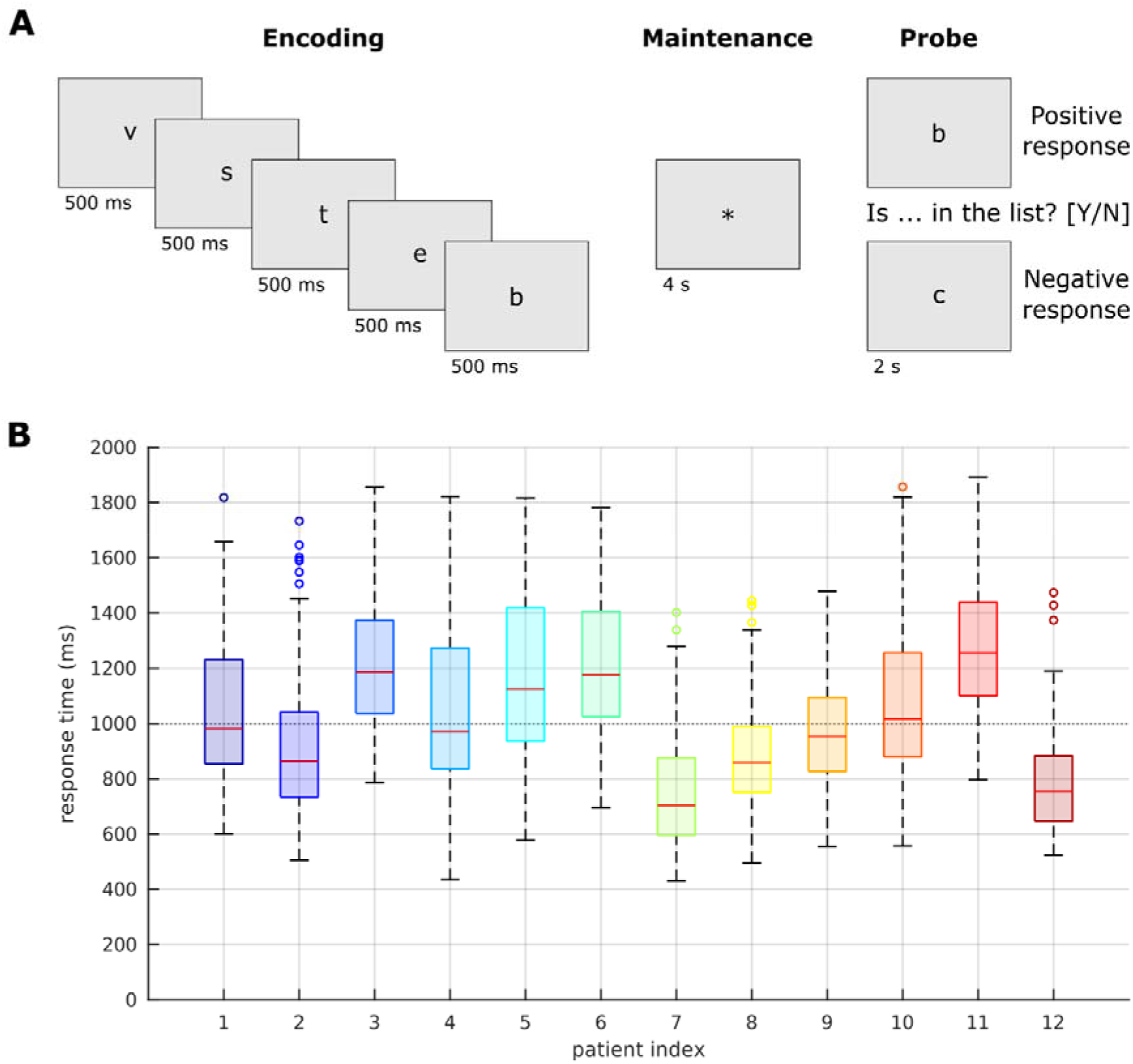
Task design and behavioral results. A. Design of the vWM task, composed of 3 phases: encoding of 5 letters, maintenance, and probe. Duration of each screen is displayed below the example frames. During the probe, the correct responses was positive (“yes, the letter was in the list”) or negative (“no, it was not in the list”). B. Response time (RT) distribution of correct trials per participant. For each patient, the red central mark represents the median RT, the bottom and top edges of the box are the 25^th^ and 75^th^ percentiles, respectively, and the lower and upper whiskers are the most extreme non-outlier RTs, respectively.

### Electrode coverage in bilateral insulae

Figure 2 shows the overall coverage of the insula. The 90 bipolar channels were distributed over the anterior and posterior regions of both insulae (35 left, 55 right; Figure 2A). In both hemispheres, the posterior long gyrus (PLG) had the largest coverage (16.7% of electrodes in the left insula and 17.8% in the right insula; Figure 2B). The main difference of coverage observed between hemispheres was the proportion of electrodes implanted in the anterior long gyrus (ALG; 4.4% for left vs. 17.8% for right).

**Figure 2:**
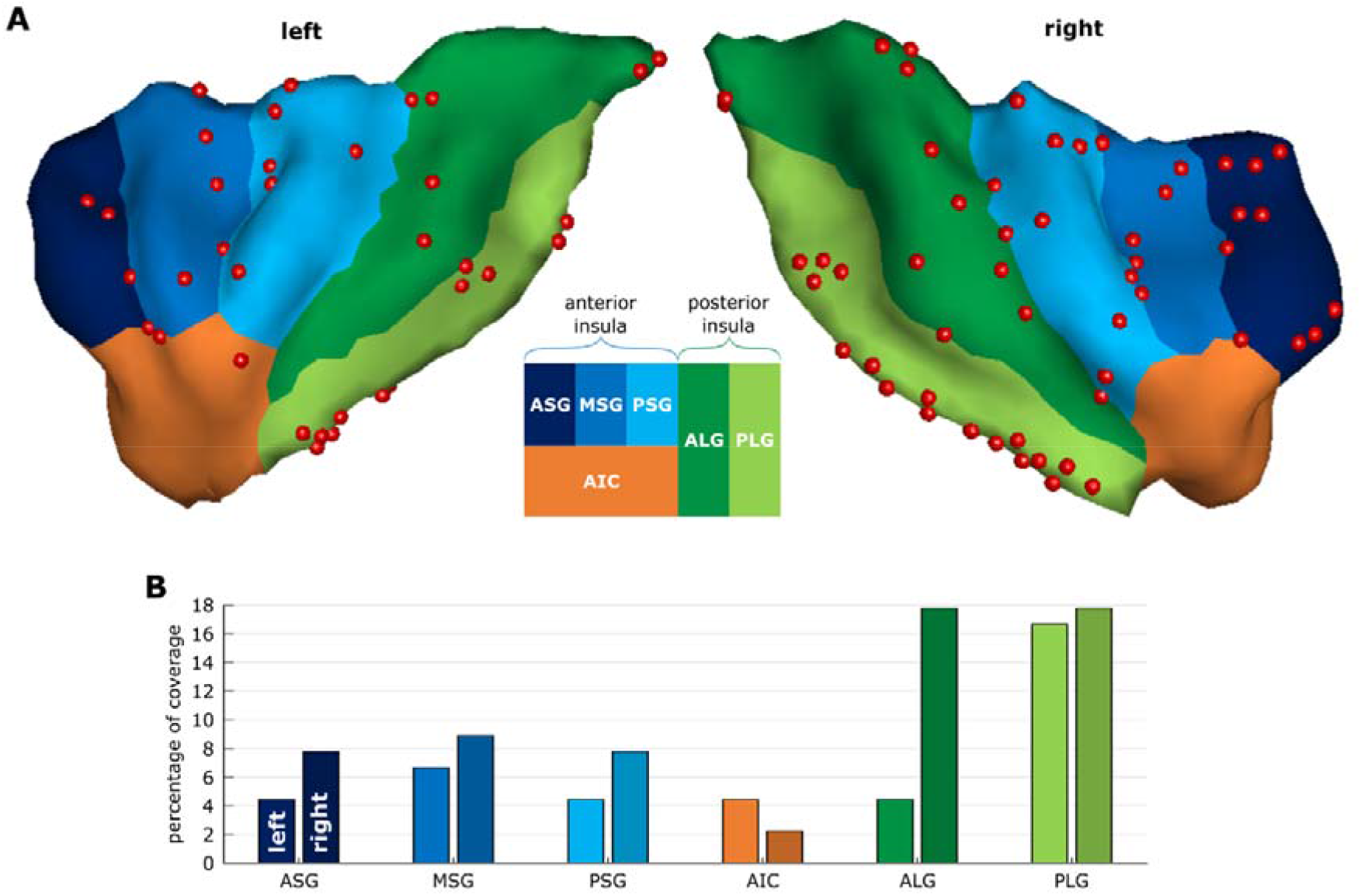
Electrode coverage for each subregion of both insulae. A. Projection of the 90 electrodes onto the 3D insula reconstructed from the MNI template brain. The color code for the 6 subregions is illustrated by the schematic representation of the different subregions. B. Percentage of electrodes in the different subregions of the left (lighter colors) and right (darker colors) insula based on patient-wise localizations. ASG: anterior short gyrus; MSG: middle short gyrus; PSG: posterior short gyrus; AIC: anterior insular cortex; ALG: anterior long gyrus; PLG: posterior long gyrus.

### Prominent frequency band modulations occurred at the probe period

A limited number of channels showed significant time-overlapping modulations (p<.05 for at least 25ms consecutively) during the encoding and maintenance periods of the correct trials for the three neural frequency bands of interest. Significant theta modulations (power changes relative to baseline) were observed in both insulae for the encoding (N = 2/12; i.e., 2 channels with significant time-overlapping modulations out of 12 responsive channels in the theta band) and the maintenance (N = 3/12) periods while beta (N = 2/9) and HFA (N = 3/26) modulations during these time periods were observed in the right insula only (see supplementary Figure). In contrast, prominent insular activity in all three frequency bands was observed at the probe period (p<.05). Based on these observations, we decided to focus the rest of the analyses on this probe time window.

In the theta band, we found that 6 of the 10 responsive channels in the left insula and 7 of the 11 channels in the right insula showed overlapping significant theta modulation during the probe period. In the left insula, the probe presentation elicited a theta increase after 400ms in electrodes implanted in its anterior part, followed by a decrease at 1400ms which was larger in the posterior insula (Figure 3A). In the right insula, a large theta increase was observed at 500ms with more channels showing a significant increase in the posterior insula, which was also followed by a second peak at 1s for 2 channels (Figure 3D). The latency difference of theta modulations between the anterior and posterior parts of both insulae was not significant (p=.38 for the left insula and p=.65 for the right insula, Figure 3G).

**Figure 3:**
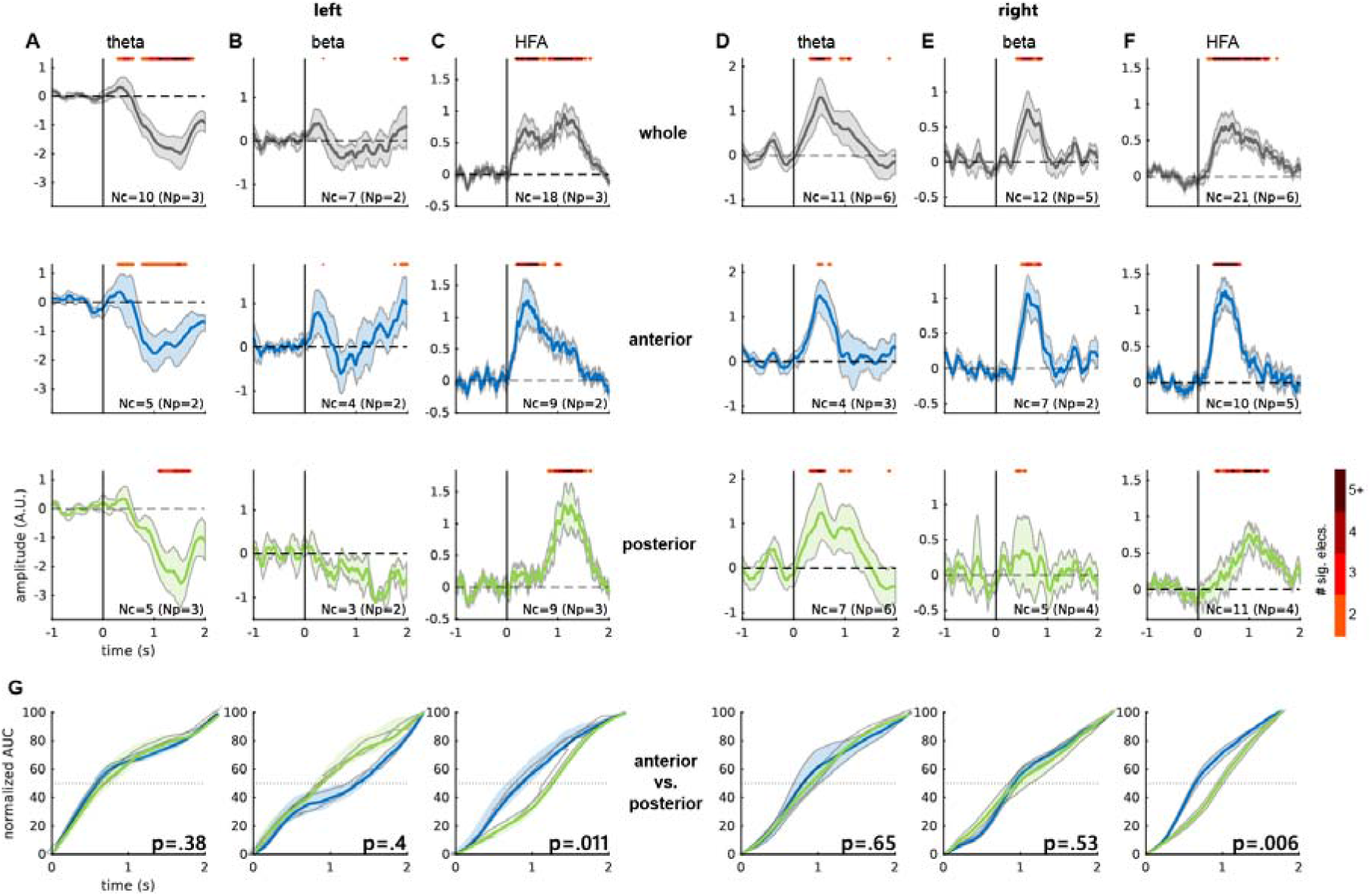
Frequency band modulations at the probe period. Time courses of theta, beta, and HFA modulations in the left (A, B, C) and right (D, E, F) insulae. Frequency band modulations averaged across all responsive channels (Nc = number of channels; Np = number of participants) for the whole insula (gray), the anterior part (blue), and the posterior part (green). Shaded areas represent the standard error of the mean (SEM). The upper red bar highlights the time points at which at least two channels showed significant modulation. The color code indicates the number of significant channels (the darker the more channels). G. Comparison of the cumulative area under the curve (AUC) for each frequency band modulations between the anterior and posterior parts (blue and green, respectively) of both insulae. The dotted line represents 50% of the normalized area under the curve.

Overlapping beta band modulations were observed in 3 out of the 7 responsive channels in the left and 4 out of the 12 channels in the right insula. In the left insula, 2 channels showed a beta increase at 300ms and 3 channels at 2s after the probe onset (Figure 3B). A significant beta increase was found around 650ms after the probe onset in the right insula, predominantly in its anterior part (Figure 3E). No timing difference was observed in the beta band along the anteroposterior axis of both insulae (left insula: p=.4; right insula: p=.5).

Finally, overlapping significant HFA modulations were observed in both insulae for 6 out of 18 responsive channels in the left and 12 out of 21 channels in the right insula. An initial increase around 400ms after the probe onset occurred in the anterior parts of the insulae bilaterally and a second HFA increase was seen 1s after the probe onset in the posterior insulae bilaterally (Figure 3C, F), with the stronger increase in the left posterior one. The latency analysis confirmed the timing difference of the HFA modulations between the anterior and the posterior parts of the insulae (p<.05 and p<.01 for the left and the right insula, respectively).

### Distinct profiles of frequency activity at the probe period

We performed a clustering analysis in order to disentangle the different profiles of frequency modulations observed in the insula. For each of the three frequency bands, we sorted the correct trials by increasing RT and formed 10 sub-averages. The optimal number of clusters of these sub-averaged time courses estimated over all responsive channels was two and differed in their pattern of frequency modulations (Figure 4A) and anatomical location (Figure 4B).

**Figure 4:**
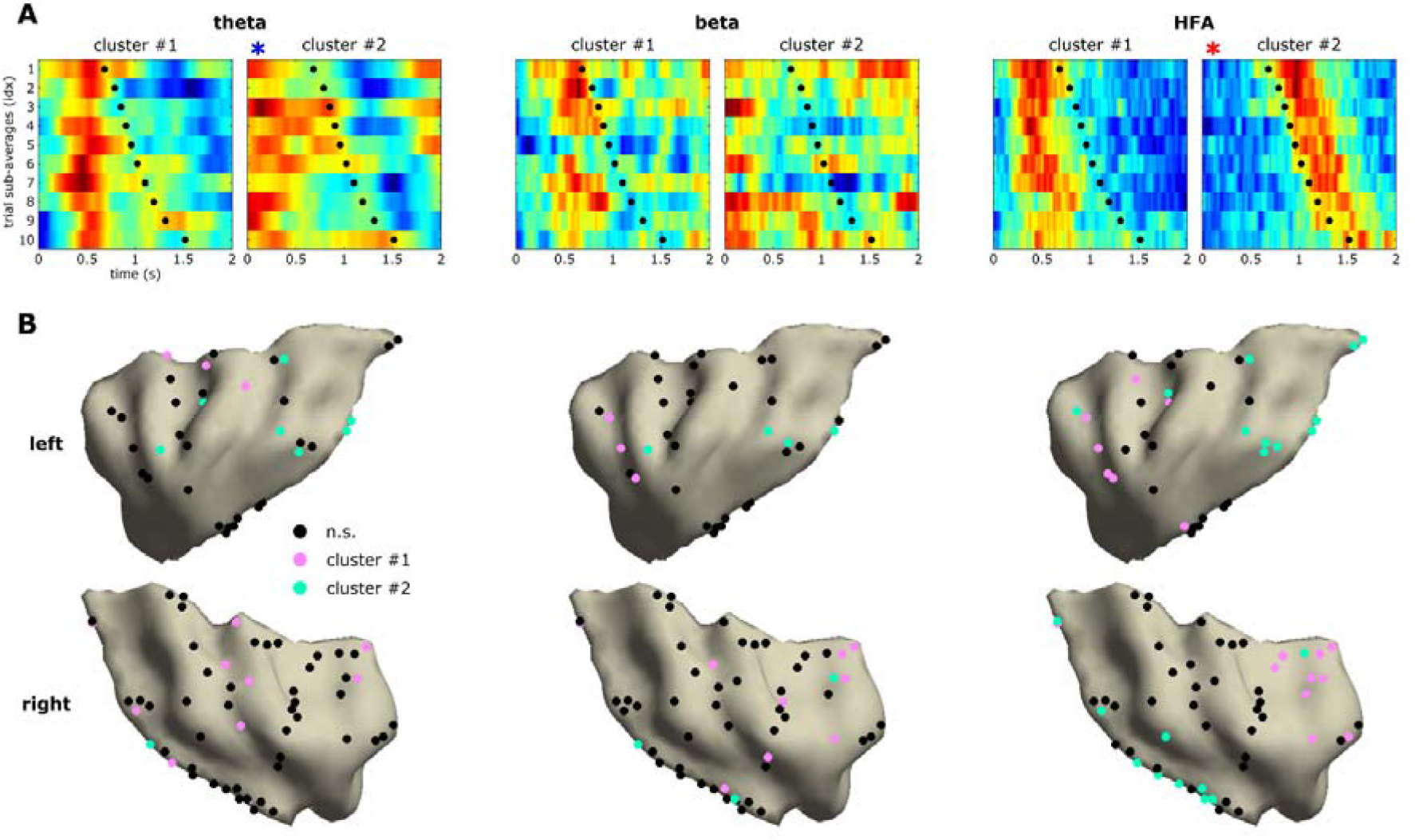
Clustering analysis at the probe period for each of the frequency bands. A. Cluster centroids as frequency band modulations sorted by increasing RT and sub-averaged, for each frequency band. Black dots represent mean RT across patients. Red represents an increase of frequency power and blue a decrease. Asterisks represent significant correlations between mean RT and peaks (red symbol) or troughs (blue symbol) of frequency band modulations. B. Cluster indices of channels per frequency band projected onto the 3D MNI insula template.

*Theta* - The first cluster was composed of 13 channels and showed a theta increase locked at 500ms after the probe onset, which was not correlated with RT (r=−.32; p=.35). This pattern of activity was predominantly found in the right insula with 10 channels (2 ASG, 1 MSG, 1 PSG, 3 ALG, 3 PLG) compared to 3 channels in the left insula (2 MSG, 1 ALG; Figure 4, left column). The second cluster revealed a theta increase at the probe onset, followed by a theta decrease with its trough correlated with RT (r=−.88; p<.001). The 8 channels showing this pattern were all located in the left insula (1 MSG, 1 PSG, 1 AIC, 1 ALG, 3 PLG), except for one (right PLG). The 7 channels in the left insula were ipsilateral to the response hand used by 2 participants of this cluster while the electrode in the right insula was contralateral to the hand used by the third participant. The spatial distribution of channels across these two clusters was different between the right and left insulae (Fisher’s exact test, p<.01) but not along the anteroposterior axis (p =1).

*Beta* - The first cluster encompassed 12 channels, predominantly in the right hemisphere (right: 3 ASG, 2 MSG, 1 AIC, 2 ALG, 1 PLG; left: 2 ASG, 1 ALG), and showed a beta increase 500ms after the probe onset. The second cluster showed a beta increase at the probe onset in 7 channels located in both insulae, with 3 channels in the anterior insula and 4 in the posterior insula (right: 1 AIC, 1 ALG, 2 PLG; left: 1 MSG, 2 PLG). None of these clusters correlated significantly with RT (r=−.68; p=.03 and r=.42; p=.23 for cluster 1 and 2 respectively; Figure 4, middle column). The spatial distribution of channels across clusters was not statistically different between hemispheres (p=.33) nor along the anteroposterior axis (p=.074).

*HFA* - The first HFA cluster encompassed 17 channels showing an HFA increase 500 ms after the probe onset, not correlated with RT (r=.39; p=.26). These channels were located predominantly in the anterior parts of both insulae (anterior insula: 8; ASG: 4 right/3 left, 7 MSG: 5 right/2left, 1 left AIC; posterior insula: 1 right ALG, 1 left PLG). The second cluster was composed of 22 channels with an increase of HFA occurring after the button press. The peak of HFA was correlated with RT (r=−.95; p<.001; Figure 4, right column). This modulation was predominantly observed in the posterior parts of both insulae (18 channels in posterior insula: 5 ALG: 2 right/3 left, 13 PLG: 8 right/5 left; 4 channels in anterior insula: 1 right ASG, 2 right PSG, 1 left ASG). These 22 channels were all located in the insula ipsilateral to the hand used by the 7 participants for whom channels were included in this cluster. The spatial distribution of channels across the HFA clusters was different along the anteroposterior axis (p<.0001) but not between insulae (p=.75).

## Discussion

Our study provides intracranial electrophysiological evidence for distributed spatiotemporal involvement of the human insula in the decision-making process required during the probe period of a vWM task. We investigated the power modulations elicited by vWM processes in three neural frequency bands within the insula. We found that the probe period elicited the strongest insula involvement, with interhemispheric and sequential intra-insular asymmetries observed across frequency bands. Significant frequency modulations in the theta band were observed during the encoding and maintenance of vWM. These results support the importance of theta modulations in successful encoding and maintenance during vWM as reported in scalp EEG^50,58^.

The probe period elicited differential involvement of the left and right insula in theta and beta bands, as well as along their anteroposterior axes in HFA. The interhemispheric asymmetry observed in our study may be due to differential roles, with the right insula being potentially more involved in the attentional aspect of the task^1,21,22^ and the left insula in the linguistic aspect^40,45^. Interestingly, a theta decrease was correlated with RT in the left insula. Theta decrease during WM has also been observed in a few electrophysiology studies using scalp EEG (see^50,58^ for reviews) and iEEG^59^. Another iEEG study showed that a theta decrease in the DLPFC was predictive of WM performance^60^. Moreover, multiple iEEG studies have reported a theta decrease, or reset, especially in the left MTL, related to successful memory retrieval^61–66^.

The time course of HFA revealed sequential activity in the insulae along an anteroposterior axis. Interestingly, limited anatomical overlapping was observed across the two HFA clusters, in line with functional differentiations between the anterior and posterior insula^21,27^. The anterior insulae were active at the probe presentation while subsequent posterior activity occurred after the behavioral response. The early HFA modulation in anterior insula provides evidence of its involvement in decision making related to the encoded verbal material^44,48^, supporting the assertion that the anterior insula is a crucial part of the cognitive control network^34,37,67–69^. The posterior insula has been reported to be involved in motor activity due to its connection with the SMA^1,2^. However, the HFA increase that correlated with response time in our study occurred after the motor response and was only found in the insula ipsilateral to the motor activity. This observation does not support the posterior insula in pure motor preparation or activation^23,29,70^. Instead, our findings support a key role in response monitoring, potentially achieved through its connection with the cingulate gyrus^7,71^. In sum, our present study implicates the insula throughout the vWM process, with a critical sequential involvement during the decision making and response monitoring phases.

### Limitations of the study

In order to confirm the role of the posterior insula in response monitoring, we ran the same pipeline of analysis over incorrect trials and no-response trials. Unfortunately, we did not find enough significant channels in any frequency band to perform a robust analysis. Further studies will be needed to better characterize the role of the posterior insula in monitoring behavior. Finally, the lack of electrodes implanted in the most ventral anterior part of insula did not allow us to make any claims on the role of this region during the vWM process.

## Materials and Methods

### Participants

Fourteen patients (4 women, mean age 32.2 years old ± 11.1 SD; 12 right-handed) with epilepsy undergoing stereotactic electroencephalography (SEEG) for pre-surgical investigation at University of California Irvine (UCI; 2) and University of Oslo (OUH; 12) performed the task. The inclusion criterion was the presence of electrodes located in the insula; 3 patients had bilateral implantation, 8 had a right implantation and 3 had a left one, for a total of 90 electrodes. All patients had an IQ score above 80. Two patients were removed from the study due to epileptic activity recorded from their insular electrodes, resulting in 12 participants. Prior to participation, each patient provided written informed consent as part of the research protocol approved by the Institutional Review Board at every site and by the Committee for Protection of Human Subjects at the University of California, Berkeley, and in accordance with the Declaration of Helsinki. The study was also approved by the Regional Committees for Medical and Health Research Ethics of Norway.

### Electrode localization

The electrodes were localized for each subject based on their individual anatomy and then transferred into the standard MNI space for representation. Affine point-based registration was used to co-register post-implantation computed tomography (CT) images to the pre-implantation magnetic resonance (MR) images using the FieldTrip toolbox^72,73^ in MATLAB (MathWorks, Inc., Natick, MA, USA).

### 3D Insula model reconstruction

To reconstruct the fine-grained 3D anatomy of the insulae, we extracted surfaces of both insulae from the MNI template brain MRI with a parcellation based on the Desikan-Killiany Atlas (aparc+aseg), using iso2mesh with 3000 vertices and 10 smoothing steps. We then manually corrected spurious vertices using a custom GUI created in MATLAB and delineated insular subregions based on two previous studies^74,75^ (Figure 2B). Finally, we snapped all electrodes onto the insular mesh at the closest vertex (Euclidean distance) of the lateral insular face (Figure 2A).

### Experimental task

The experimental paradigm was a Recent-Probes task^76,77^ (Figure 1A). Each trial was composed of a list of five letters displayed on the screen for 500ms each, with an interstimulus interval of 500ms. This encoding phase was followed by a 4-second period of maintenance (represented by a star on the screen). A question was then displayed for 2s asking whether a given letter, i.e., the probe, was in the current list. The probe letter was present in the current list for half of the trials (positive condition) and absent from it for the other half (negative condition). The task included a total of 144 trials in pseudorandomized order for each participant. The trials were presented in three blocks of 10 min each, with short breaks between blocks, resulting in a total study duration of approximately 35 min.

### Experimental setup

The participants were seated on their hospital bed, with the laptop placed in front of them on an overbed table. A photodiode was placed on the bottom left corner of the laptop screen and recorded as a channel (UCI), or alternatively a digital trigger (TTL) was used to mark each timestamp of interest (OUH).

The instructions were displayed on the screen and required the participants to respond ‘‘yes’’ (positive response condition) or ‘‘no’’ (negative response condition) by pressing the mouse keys using their index and middle finger, respectively. To prevent potential interference of motor activity due to button presses occurring simultaneously with the electrophysiological signals of interest, the hand used for the task was ipsilateral to the implantation when the implantation was unilateral. For bilateral implantations, one participant used their right hand while the two other participants used their left hand.

### Behavioral analysis

The response time at the probe was recorded for each trial. Trials with RT faster than 400ms^77^ or exceeding three standard deviations were excluded^78^. The remaining trials were included in subsequent analyses of response latency (based on the mean RT of correct trials per participant) and error rate (based on the percentage of incorrect trials). We performed Wilcoxon signed-rank tests in MATLAB to test the hypothesis of similar mean RT between positive and negative answers for correct trials, and to assess a potential difference in error rate between positive and negative answers.

### SEEG data acquisition

Two datasets were acquired at UCI using a Nihon Kohden recording system (Nihon Kohden Corporation, Japan) sampling at 5kHz. The 10 other datasets were recorded at OUH: 4 datasets were acquired using a NicoletOne system (Nicolet, Natus Neurology Inc., USA), 1 dataset using a 256-channel Brain Quick system (Micromed, Italy), and 5 with an ATLAS system (Neuralynx, USA). The sampling rates were 512Hz, 2048Hz, and 16kHz respectively. Each SEEG shaft was composed of 5 to 18 electrodes spaced 5mm apart for UCI (Ad-Tech, Racine, WI, USA) and 3.5mm apart for OUH (DIXI Medical, France).

### SEEG data preprocessing

Raw data with a sampling rate higher than 1kHz was downsampled offline to 1kHz and filtered with 1Hz high-pass and 250Hz low-pass filter (4^th^ order Butterworth). In addition, 60Hz for UCI and 50Hz for OUH power-line noise frequency and harmonics were removed using notch filters. The filtered data was then visually inspected under the supervision of a neurologist (RTK) to identify and discard from further analyses channels and epochs displaying epileptiform activity or artifactual signals (from poor or broken contact, machine noise, etc.). Clean, artifact-free, continuous data was epoched into 16s-long trials (from −12 to 4s, 0s being the probe presentation; epochs included 2s of data padding at each extremity to avoid edge artifacts in the following time-frequency (TF) decomposition).

Each trial encompassed the encoding of the five successively presented letters (from 9 to 4s before the probe presentation), the maintenance of the letters (4s leading up to the probe presentation at 0s), and the probe period (from 0 to 2s post-probe onset). Behavioral information of each trial (condition, RT, and accuracy) was preserved during the epoching. The epoched data was re-referenced using bipolar montage (i.e., subtraction of the activity of a given electrode from the previous neighbor electrode on the same SEEG shaft^79^, yielding 96 virtual channels with minimized contamination from volume conduction^80–82^. A visual re-inspection of the bipolar data was performed to reject any remaining channels or epochs containing residual artifacts. The complete preprocessing yielded a total of 90 artifact-free bipolar channels (with at least one electrode in the insula for each channel). All preprocessing steps and subsequent analyses were performed using the FieldTrip toolbox for MATLAB and custom MATLAB scripts.

### Time-Frequency decomposition

We performed a TF decomposition of all correct trials using Morlet wavelets, over a frequency range from 4 to 150Hz (logarithmically spaced in 32 steps) and using a time resolution of 5ms resulting in a sampling rate of 200Hz. The wavelet order (width or number of cycles) was logarithmically increased from 4 to 30 cycles (also in 32 steps). A baseline correction (with the second preceding the encoding onset used as baseline period) was then applied using the decibel method ^2^. Lastly, the padding time periods were removed so that the remaining epochs lasted 12s (−10 to 2s relative to the probe presentation).

### Analyses over the entire vWM process

We first extracted the time courses of the 3 frequency bands of interest (theta: 4-8Hz; beta: 13-30Hz; High-Frequency Activity, or HFA: 70-150Hz), by averaging the corresponding baseline-corrected frequency bins of the TF decomposition. For each frequency band and each channel, we computed a paired-sample t-test comparing each time sample contained in the trial (−9 to 2s) against the baseline mean (−10 to −9s). We then isolated the significant time samples that stayed significant (p<.05) for at least 25 consecutive ms using a Bonferroni correction to compensate for multiple comparisons (alpha value divided by the number of time samples) and labeled the channel as responsive. Frequency modulations were considered significant if at least 2 responsive channels were significant for the same time sample, regardless of the patient (see supplementary Figure).

### Analyses at the probe period

Similar parameters (i.e., p<.05, 25 consecutive ms) as for the entire time window were used to extract the frequency band time courses at the probe period (0 to 2s]). The paired sample t-tests to select the responsive electrodes were computed against the mean of the second preceding the probe onset (−1 0s).

We assessed the difference of frequency band modulation latencies between the anterior and posterior parts of the insula by using the fractional area method ^84^. For each frequency band and each responsive channel, we first set the minimum of the time course to zero and normalized it between 0 and 100%. We then computed the cumulative area under the curve (AUC) over the time window of interest (0 to 2s). The time point separating the cumulative AUC in two equal parts (50%) was extracted. A Wilcoxon rank sum test was performed to assess latency differences between anterior and posterior insular channels.

### Spatiotemporal k-means clustering analyses at the probe period

For each responsive channel, correct trials were sorted by ascending RT and then grouped into 10 bins of trials equitably distributed across bins (for denoising purposes). Note that due to varying numbers of correct trials across patients, the tenth bin sometimes contained less trials than the nine other bins. We then extracted the mean RT and the mean temporal profile of each bin of the probe time window (0 to 2s), resulting in a 10-by-400 matrix (2s epoch at a 200Hz sampling rate).

We then performed a k-means analysis to cluster these RT-sorted temporal profile matrices, based on the nearest centroid using the Squared Euclidean distance^85^. For each of the 3 frequency bands of interest, we evaluated the optimal number of clusters using the Calinski-Harabasz clustering evaluation criterion, replicated 500 times. This optimal number analysis revealed the presence of two main clusters for each frequency band. We then visualized cluster centroids and plotted the channels of each cluster on both insulae.

We compared the spatial distribution of the channels within the k-means clusters obtained for each frequency band by performing Fisher’s exact tests. For each frequency band, we compared antero-posterior and inter-hemispheric distributions of channels across clusters.

### Correlation with behavioral response time

For each frequency band and for each cluster, we performed linear correlations between the peak (or trough) of activity of the cluster centroids, i.e., the latency of the maximum (or minimum) value of each of the 10 sub-averaged temporal-profile bins, and the mean RT. A Bonferroni correction was performed to compensate for multiple comparisons by dividing the alpha value by the number of correlations (number of clusters by number of frequency bands).

## Acknowledgments

We acknowledge the members of the Oslo team and the Knight Lab, especially Dr. Colin Hoy, for their help during the analysis and the discussion of the study, and the patients who participated in the study. The authors also want to acknowledge Pr. Ulrike Kraemer who helped us initiate the project. This study was supported by NINDS R37NS21135 and Brain Initiative U19 U19NS107609 (to RTK), the Research Council of Norway (project number 240389 to A-KS and AOB, 314925 to AOB, and 274996 to A-KS and TE), and the Research Council of Norway through its Centres of Excellence scheme (project number 262762, RITMO).

## Authors contributions

Conceptualization: TE, AKS, AL, LB

Methodology and Analysis: LB, AL

Investigation: AL, LB, AOB

Resources: JI, PGL, JJL

Writing Original: AL, LB, RTK

Writing review and Editing: AOB, JI, PGL, JJL, TE, AKS

Supervision and funding acquisition: TE, AKS, RTK

## Data and code availability statement

The datasets generated and analyzed during the current study are available from the corresponding author upon reasonable request. All the scripts used for the analyses are available online at: https://github.com/ludovicbellier/InsulaWM

## Ethics Statement

Electrode placement was solely determined based on clinical considerations and all procedures were approved by the institutional review boards at the hospitals, as well as the University of California, Berkeley and by the Regional Committee for Medical Research Ethics, Region South Norway, and in accordance with the Declaration of Helsinki. The participants provided their written informed consent to participate in this study.

## Supplementary information

**Supplementary Figure:**
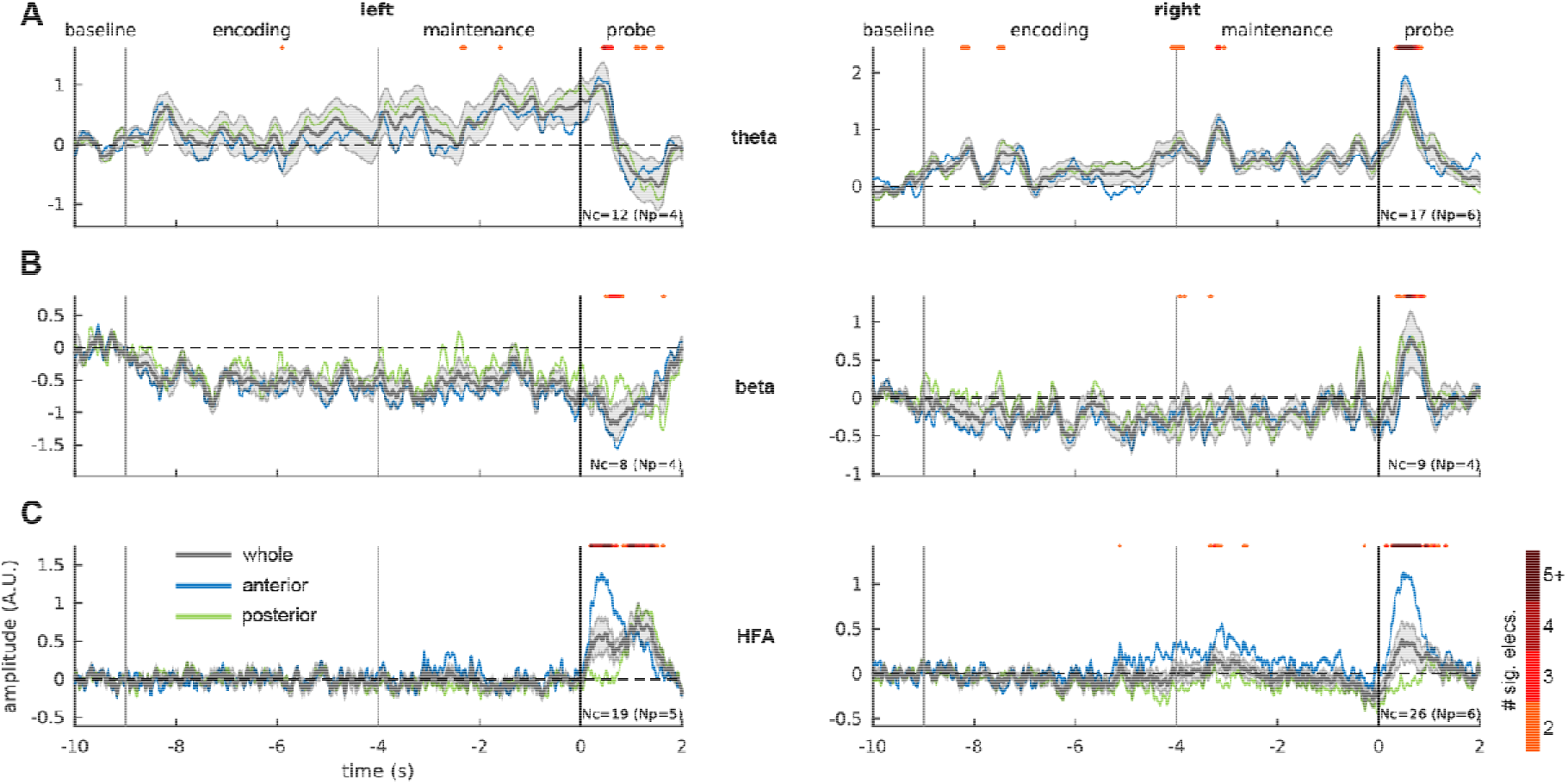
Frequency band modulations during the entire vWM process. Time courses of theta (A), beta (B), and HFA (C) modulations in the left and right insulae. Frequency band modulations averaged across all responsive channels (Nc = number of channels; Np = number of participants) for the whole insula (gray), the anterior part (blue), and the posterior part (green). Shaded areas represent the standard error of the mean (SEM). The upper red bar highlights the time points at which at least two channels showed significant modulation. The color code indicates the number of significant channels (the darker the more channels).

1 The gender and ethnic proportions of our reference list has been checked using CleanBib (https://github.com/dalejn/cleanBib). The reference list contains 7% woman(first author)/woman(last author), 14.4% man/woman, 26.8% woman/man, and 51.8% man/man; and 12% author of color (first)/author of color(last), 13.4% white author/author of color, 20% author of color/white author, and 54.6% white author/white author. These proportions are in the expected range of gender and ethnicity ratios currently present in the field of neuroscience^86–88^; not included in the reported statistics). Note that this classification is based on names, pronouns and online databases and does not account for intersex, non-binary, transgender, Indigenous and mixed-race authors.

